# Failure properties and microstructure of porcine aortic adventitia: fiber level damage vs tissue failure

**DOI:** 10.1101/2023.03.13.531658

**Authors:** Venkat Ayyalasomayajula, Baptiste Pierrat, Pierre Badel

**Affiliations:** Department of Structural Engineering, Norwegian University of Science and Technology, Trondheim, Norway; Mines Saint-Etienne Univ Lyon Univ Jean Monnet INSERM U 1059 Sainbiose Centre CIS F - 42023 Saint-Etienne France

**Keywords:** aortic rupture, collagen fibers, damage modeling

## Abstract

Aortic aneurysm rupture is a sudden local event with high mortality. It is generally accepted that the adventitia acts as the final barrier protecting the aorta from over-expansion. Currently, the knowledge of microscopic structural determinants of the tissue’s mechanical response and failure is very limited. The purpose of this study is to provide data on the directional failure properties of the adventitia, combined with micro-structural imaging and structure based constitutive modeling to quantify fiber-scale rupture criteria. Eleven healthy porcine aortas were used in this study. Cylindrical portions of the abdominal section were excised, cut-open longitudinally, the medial and adventitial layers separated methodically. Picrosirius red staining was used to image the collagen fiber morphology via an optical microscope. Subsequently, dog-bone shaped specimens were subjected to uniaxial testing until failure while being recorded by a Nikon digital camera. A fiber-scale damage model was utilized to explain the tissue-scale failure. The ultimate tensile stress in the circumferential and longitudinal directions were recorded to be 0.96 ± 0.29 *MPa* and 0.85 ± 0.36 *MPa* respectively. Meanwhile, the ultimate stretch to failure in the circumferential and longitudinal directions were recorded to be 1.72 ± 0.16 and 1.88 ± 0.13 respectively. Further, correlation between the failure properties of the tissue and mean fiber orientation have been reported. Finally, the critical fiber stretch for damage initiation and eventual tissue failure were identified to be 1.19 ± 0.07 and 1.24 ± 0.05 for circumferential and longitudinal specimens respectively. Our approach provides valuable insight into the (patho)physiological mechanical role of collagen fibers at different loading states. This study is useful in enhancing the utilization of structurally motivated material models for predicting arterial tissue failure.

## 1. Introduction

The aorta is the largest artery in the human body, which carries oxygenated blood from the heart to the rest of the body. Among the many conditions that affect the aorta, aortic aneurysms (AA) are often considered the most dangerous and difficult to treat with high mortality rates [1, 2]. An aneurysm within an aorta is identified as a focal dilation, with a diameter greater than 30mm. Current clinical practice guidelines are: aneurysms with a diameter of 30 - 39 mm, medical therapy, and watchful waiting is recommended. For medium-sized aneurysms (40 - 54 mm), checkups every 4-6 months are indicated, whereas AAs that are 55 mm or more in diameter, surgical intervention is recommended [3, 4]. AA diameter remains the most widely used criterion for intervention, which is based in part on a retrospective review of 24000 autopsies over a period of 23 years. However, from the above study, 40% of AAs with a diameter between 70–100mm were unruptured, whereas 13% of AAs with a diameter less than 50mm were ruptured [5, 6]. Clearly, this “one size fits all” criterion fails many individuals and is not adequate in the era of personalized medicine. Consequently, better knowledge of the mechanics of aortic tissue failure can be important for predicting adverse events such as aneurysm rupture.

The healthy aorta is a composite structure, made up of three distinct concentric layers: tunica intima, tunica media, and tunica adventitia. All the arterial walls are composed of connective tissue structures: endothelial cells, smooth muscle cells, collagen, and elastin fibers; arranged in these three layers. Within a healthy aorta, the medial layer, composed mainly of smooth muscle cells, elastin fibers, and some collagen is responsible for the main structural and functional properties of the aorta [7, 8]. The outermost adventitial layer is made of thick bundles of undulated collagen fibers in the unloaded state making it very compliant at small strains but changes to a stiff tube at high strains thus preventing the artery from over-stretching and rupture [9, 10]. Within a diseased aorta (such as an aneurysmal aorta), a key difference is the degeneration of the medial layer: typically involving the disintegration of smooth muscle cells and fragmentation of elastin fibers [11] and an increase in collagen content [12]. The three distinguishable layers of a normal aorta is reconstituted to a single layer akin to the adventitial layer of the healthy aorta with an abundance of collagen fibers and a fragmented network of elastin fibers [13]. Further, the collagen fibers in the diseased aortic wall tend to lose their waviness and appear as thick straight wires [14]. Elastic arterial biomechanics has been intensively studied. Mechanical testing protocols such as inflation tests [15, 16], simple uniaxial tension [17, 18], and biaxial tests [19, 20] have been used to determine the elastic mechanical properties of aortic tissues. Further, there are many reported experimental analyses of failure mechanics of arteries, which determine the peak strength of the tissue [21, 22, 23]. However, only a limited number of studies report on the layer-specific strength of the tissue, which are often limited to the tunica media [24, 25]. The role of adventitia as the final barrier is often overlooked. Only a handful of studies have reported the mechanical strength of and failure properties of the adventitial layer [26, 27]. Further, there are very few experimental investigations of the damage and rupture mechanisms of arteries [28, 29]. Weisbecker et al [18] addressed the layer-specific softening response of the human aorta. Yet, their study only presented the softening response of the layers and there were no data about the failure and rupture properties. A few studies [30, 24] combined micro-structural imaging with mechanical testing, however, they do not provide any explanation of damage mechanisms in the arterial tissue. Clearly, damage mechanisms and specific injury tolerance are closely related to the organization and content of micro-structural constituents in the tissue. Specifically, collagen seems to play a universal pivotal role. Therefore, understanding and quantifying damage mechanisms at the scale of collagen fibers is important to be well characterized.

Several damage formulations have been proposed to describe the failure of arteries [31, 32, 33]. A detailed review of damage models for soft biological tissues is presented in [34]. The mechanical properties of the collagen fibers and their contribution to the overall behavior of the tissue being the main load-bearing constituents of soft tissues is of particular interest. As reported by Pins and Silver [35], softening and rupture of collagen fibers is a phenomenon that happens under supra-physiological loading or in the case of disease. But as yet very few studies focus on incorporating such micro-scale effects in constitutive models. For instance, the brittle failure mechanism for collagen fibers assumes that fibers fail abruptly once a critical stretch is reached [36, 37]. It reproduces the typical non-linear behavior, including the toe and linear regions, then damage and eventual failure. Gasser proposed an irreversible damage model arising from the scale of proteoglycan bridges to account for plastic deformations [38]. Alastrue [39] proposed a macroscopic tissue damage model by considering recruitment stretches for the fiber content and failure of fibers in distribution at different stretches or strain energies. Decoupled damage mechanisms for the matrix and fibers are considered. More recently, Holzapfel et al [40] proposed a model to account for progressive damage in collagen fibers and incorporated it into their well-established structure-based strain energy function [41].

The purpose of this study was to characterize the failure properties of the adventitial layer of the healthy descending abdominal aorta combined with micro-structural imaging in order to quantify a basic model at the collagen fiber level that can account for softening and rupture. To achieve this, we performed uniaxial tests to study the mechanical behavior of layer-separated porcine aortic tissue samples taken from descending abdominal aorta until complete failure. For its simplicity and applicability, a structure-based strain energy law [14] and its extension to a micro-structural damage model [40] are considered in this work to reproduce the experimental data. The choice of the damage model was made following observations made during performing the uniaxial tensile tests under a Nikon Digital camera where fiber-sliding and fiber-bundle rupture were apparent leading to tissue failure. Finally, we evaluate structure-function relationships to study the influence of collagen morphology on the failure properties of the tissue.

## 2. Methods

### 2.1. Sample preparation

Healthy porcine aortas (n = 11) in whole were procured from a local butchery within 24 hours of death. Once in the lab, the aortas were cleansed with ionized water and the surrounding connective tissue was carefully removed. Cylindrical segments of lengths 5 cms were excised around the descending portion of the aorta and cut open longitudinally. This resulted in rectangular samples of length 5 cms and width 3 – 4*cms* (see figure 1). The medial layer of the aorta was carefully peeled off from each excised sample resulting in adventitial strips. These resulting adventitial strips were frozen at −20°*C* until the day of the experimental testing. On the day of the experiment the frozen adventitial samples excised from healthy porcine aortas were unfrozen in ionized water. Dog-bone shaped samples were cut in both circumferential and longitudinal alignments for mechanical testing.

**Figure 1:**
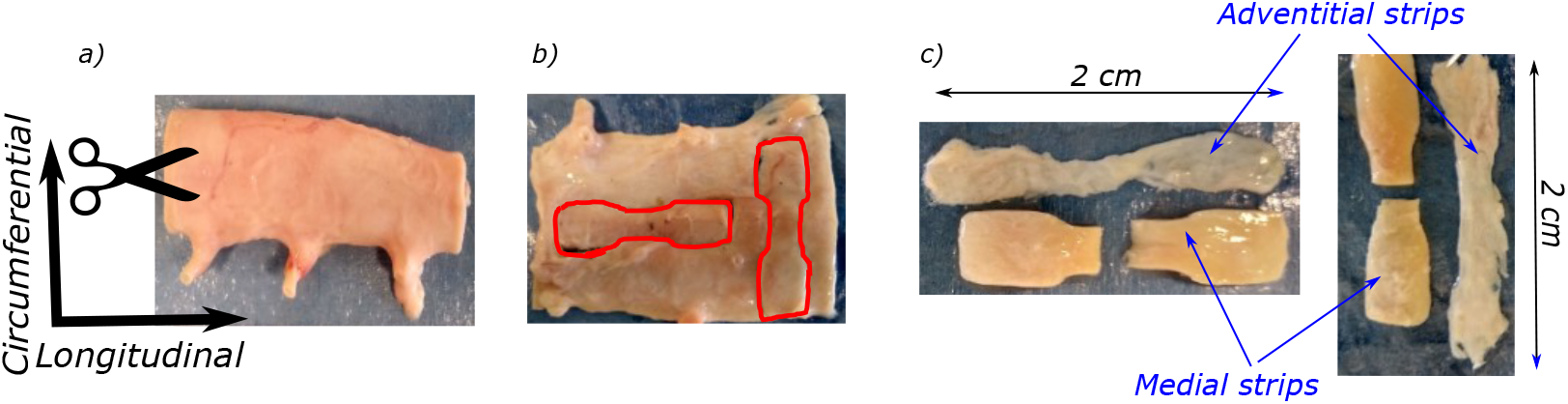
Whole porcine aorta used for mechanical testing, b) schema of prepared dogbone specimens, c) excised dog-bone specimens with the medial layer separated before testing.

### 2.2. Thickness measurement protocol

The thickness measurement device [23] consisted of two high spatial resolution line laser sensors (optoNCDT 1700BL, Micro-Epsilon Messtechnik GmbH & Co. KG, Germany) facing each other in a horizontal line and a central body midway between the sensors. The central body consisted of a linear translation stage (MFA-PPD, NewPort, CA, USA) capable of moving the sample in the vertical direction. Two PVC supports were used to hold the sample in between them, which were then attached to the movable translation stage. Two laser beams in the region of blue light (λ = 405nm) were chosen such that soft tissues reflected these wavelengths without significant light absorption and blur, as demonstrated for skin [42]. These incident laser beams were used to scan the surface of the tissue on either side. The scanner exposure time was 1 ms and the total measurement field was 1280 × 768 pixels. A program developed in LabVIEW (National Instruments, Austin, USA) was used to record the measured position profiles of both exposed surfaces of the sample with a vertical step of 50 *μm* and a lateral resolution of 20 μm. Measurements on porcine aortic adventitial strips was made once with a total scan time of about 2 minutes. A custom Matlab program was used to determine the thickness of the tissue. This was done by first mapping the two surfaces on to each other, which allowed for defining a vector normal to the mean surface. Finally, the thickness was computed as the distance between the two surfaces along this normal. Thickness profile of a representative human porcine aorta (only the adventitia) is presented in figure 2.a.

**Figure 2:**
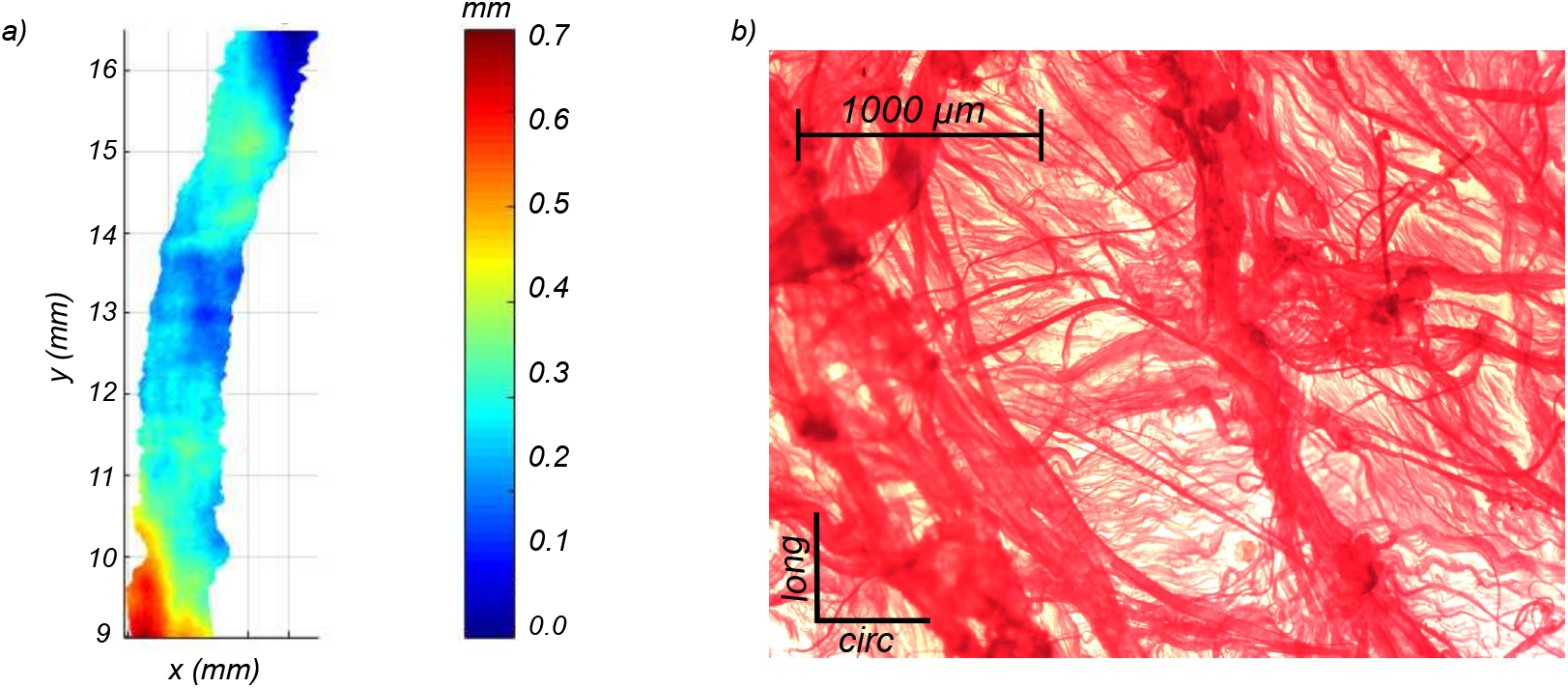
a) The result of a thickness scan giving a map of thickness measurements for the whole specimen, b) Adventitial collagen fiber architecture imaged under Optical light microscope at a magnification of 4×.

### 2.3. Optical microscopy

For optical light microscopy, a fluorescence compatible LEICA DM750 P educational microscope equipped with standard LED illumination was used. The cylindrical samples were unfrozen in ionized water for 15 – 30 mins and cut open longitudinally. The prepared adventitial samples were stained in picro-sirius red solution for one hour and washed in ionized water. At this point, the prepared samples were placed in-between two glass slides in-order to protect the sample and the objective lens and mounted on to the stage. Images of collagen architecture as shown in figure 2.b were procured at two different magnifications: 4× and 10× giving insights into multi-scale (fiber bundles and fibers) organization of collagen fibers in the adventitia.

### 2.4. Uniaxial testing

The mechanical testing was conducted using a screw driven high precision tensile machine (Newport®, tension-compression stage) with a load cell capacity of 22 N. The load cell was integrated to an IPM650 panel mount for digital display of load cell readings. Each sample was carefully mounted on to the testing machine and clamped at the ends with the help of sandpaper so as to prevent slipping. It was also ensured that the sample was submerged under water for the entirety of the test. Ensuring a zero-strain state in the initial configuration before preconditioning was achieved by maximally constraining and controlling sample and clamp positions, upon fixation (figure 3). The zero-stress initial configuration was achieved by resetting to 0 the load cell offset before mounting each sample, and by verifying the unchanged force readout after mounting the sample. This verification gave an a posteriori check that the fixation did not induce any pre-stress of the sample larger than the force threshold being detected by the sensor (i.e. 0.02 N, which corresponds to 30 kPa according to our sample’s mean cross-sectional area). A displacement control loading was applied to each sample at a rate of 2 *mm.min*^-1^ until failure.

**Figure 3:**
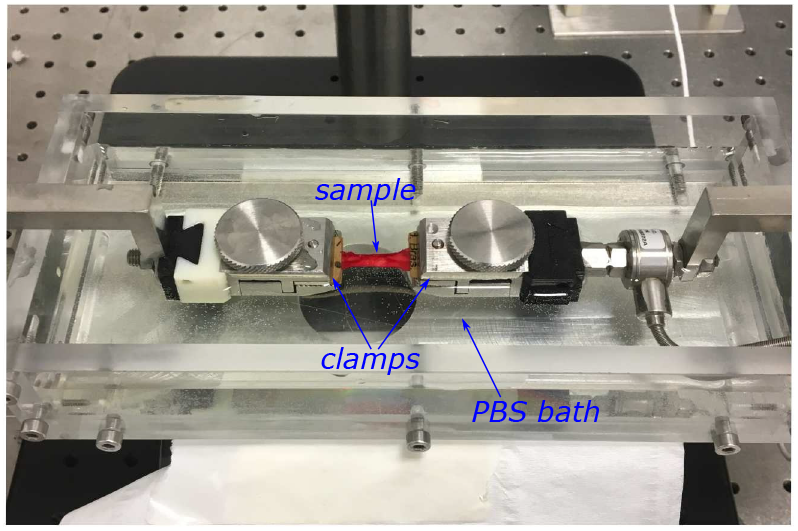
Uniaxial tension set up used for testing porcine aortic tissue samples.

Each tested sample followed a preconditioning protocol, where 5 cycles of a triangular load profile of up to 30% strain and unloading was applied. The above mentioned zero stress and zero strain conditions are related to the initial configuration before preconditioning. It was decided not to account for any residual strain from preconditioning when plotting the stress-strain curves of the samples. After preconditioning, three measurements of the sample length and width in the unloaded configuration were recorded using a caliper and averaged. This measured sample width in conjunction with the mapped thickness profile of the sample allowed for the computation of the reference cross-sectional area *A*_0_ (at zero load, after preconditioning). From this, the first Piola-Kirchoff stress was computed as: 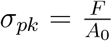, where *F* is the measured tensile force. The knowledge of initial inter-clamp length allowed for the computation of stretch as: 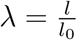 where *l* and *l*_0_ are current and initial inter-clamp lengths respectively.

A Nikon 5600 digital camera was used to image the sample through out the test at each intermediate loading state. The set-up was aimed at imaging the fibers and capture their reorientation with progressive tensile load. To achieve this the samples were stained with Picrosirius red for 30 mins prior to mechanical testing. The recorded images for a circumferential and a longitudinal sample at their initial configuration and close to failure is presented in figure 4 a-b and figure 4 c-d respectively.

**Figure 4:**
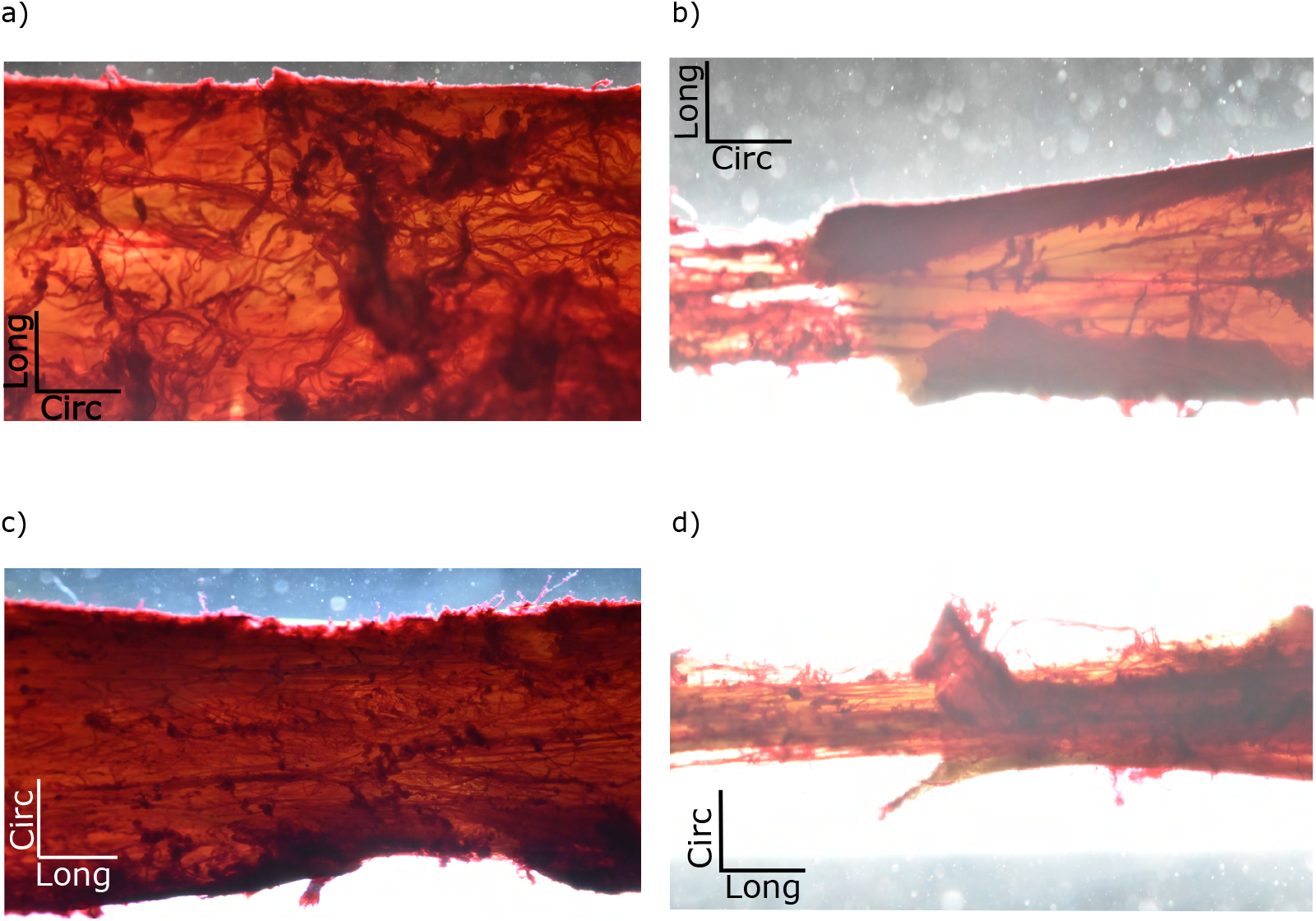
Circumferntial sample at load-free state, b) Circumferential sample at close to failure under uniaxial tension, c) Longitudinal sample at load-free state, d) Longitudinal sample at close to failure under uniaxial tension.

### 2.5. Damage model

The stretch of the material *λ* in any given direction represented by a unit vector e is defined as the ratio between its length in the deformed configuration and in the reference one, expressed as: *λ*^2^ = *e*: *C*: *e*. Here, *C* = *F^T^F* is the right Cauchy-Green tensor, and *F* = *∂x/∂X* represents the deformation gradient tensor.

The pseudo-elastic damage model developed by Holzpafel et al in [40] is implemented here without the inclusion of cross-links. Briefly, the elastic damage model postulated in terms of the strain energy density is given as:

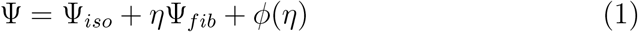

where,

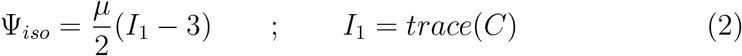

and *μ* > 0 is a material parameter and *I*_1_ is the first invariant of C. Similarly, the anisotropic contribution of the fibers is defined as:

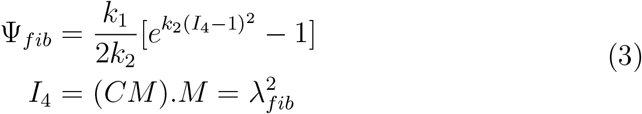

where, *M* = *cos*(*θ_m_*)*e_x_* + *sin*(*θ_m_*)*e_y_* is the unit vector along mean fiber orientation of the fibers in the load-free configuration and *λ_fib_* is the fiber stretch. The variable *θ_m_* represents the mean orientation angle with *e_x_* and *e_y_* being unit vectors in uniaxial loading direction and its orthogonal respectively. The damage variable and damage function are given by:

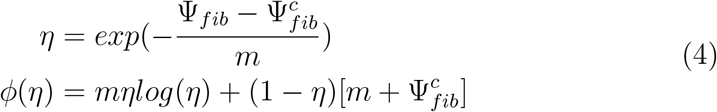

Here *m* > 0 is a material parameter and 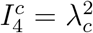 is the critical stretch value in the fiber direction, responsible for the initiation of damage. As *η* decreases from 1 to 0 for *λ_fib_* > *λ_c_*, the tissue approaches global failure.

For the case of uniaxial tension, the analytical expression for stress derives to:

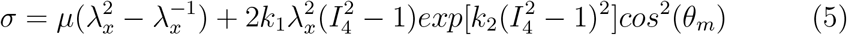

when there is no damage in the tissue and for *λ* > *λ_c_* we obtain:

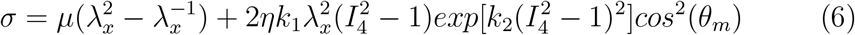

There are in total 6 parameters in the model including *μ, k*_1_, *k*_2_, *m, λ_c_*, and *θ_m_*. *θ_m_* is determined from optical microscopy images. The isotropic material parameter *μ* is estimated inversely by making comparisons to experimental stress-stretch curves within the low-strain regime (*λ* < 1.05). This is done taking into consideration that collagen fibers get recruited into principle load bearing with stretch. The anisotropic material constants *k*_1_ and *k*_2_ are identified on the elastic part of the stress-stretch response. Finally, the damage/failure parameters *m* and *λ_c_* are identified on the damage part of the stress-stretch response.

## 3. Results

A total of 11 porcine abdominal aortic sections were examined, with one circumferential and one longitudinal specimen excised from each aorta.

### 3.1. Mechanical testing

Uniaxial tension tests were performed on the excised specimens until failure according to the experimental procedure described in section 2.4. In this case, the the accurate micro-scale rupture mechanisms could not be identified and formulated. Although a mixture of de-cohesion between fiber bundles and individual fiber rupture were observed (see video attached in supplementary material). However, the tissue’s aggregate elastic and failure mechanical properties are identified. The recorded stress-stretch curves until failure for all specimens is presented in figure 5. The average thickness of the adventitial layer was measured to be 328 ± 47 μm. Table 1 presents the overall recorded mechanical response parameters for all specimens. With reference to the failure stress, the circumferential specimens had a higher strength (0.96 ± 0.29 *MPa*) as compared to the longitudinal specimens (0.86 ± 0.35 *MPa*). However, t-test confirms that the two groups were not statistically different (*p* – *value* – 0.0184). On the contrary, longitudinal specimens had a higher stretch to failure (1.88 ± 0.13) as compared to circumferential specimens (1.72 ± 0.16). Again, the two groups were not statistically different (*p* – *value* – 0.0142). The descriptive statistics of the mechanics of all specimens are shown in figure 7.

**Figure 5:**
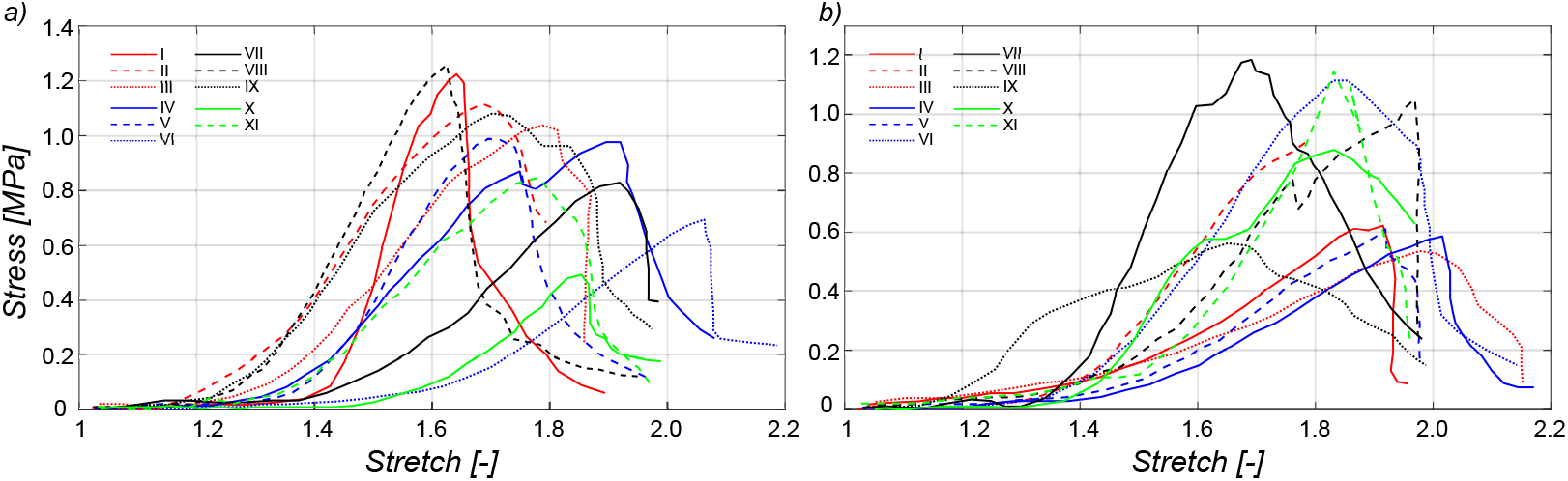
Uniaxial tension stress-stretch curves until failure for all porcine adventitial specimens: a) circumferential, b) longitudinal.

**Figure 6:**
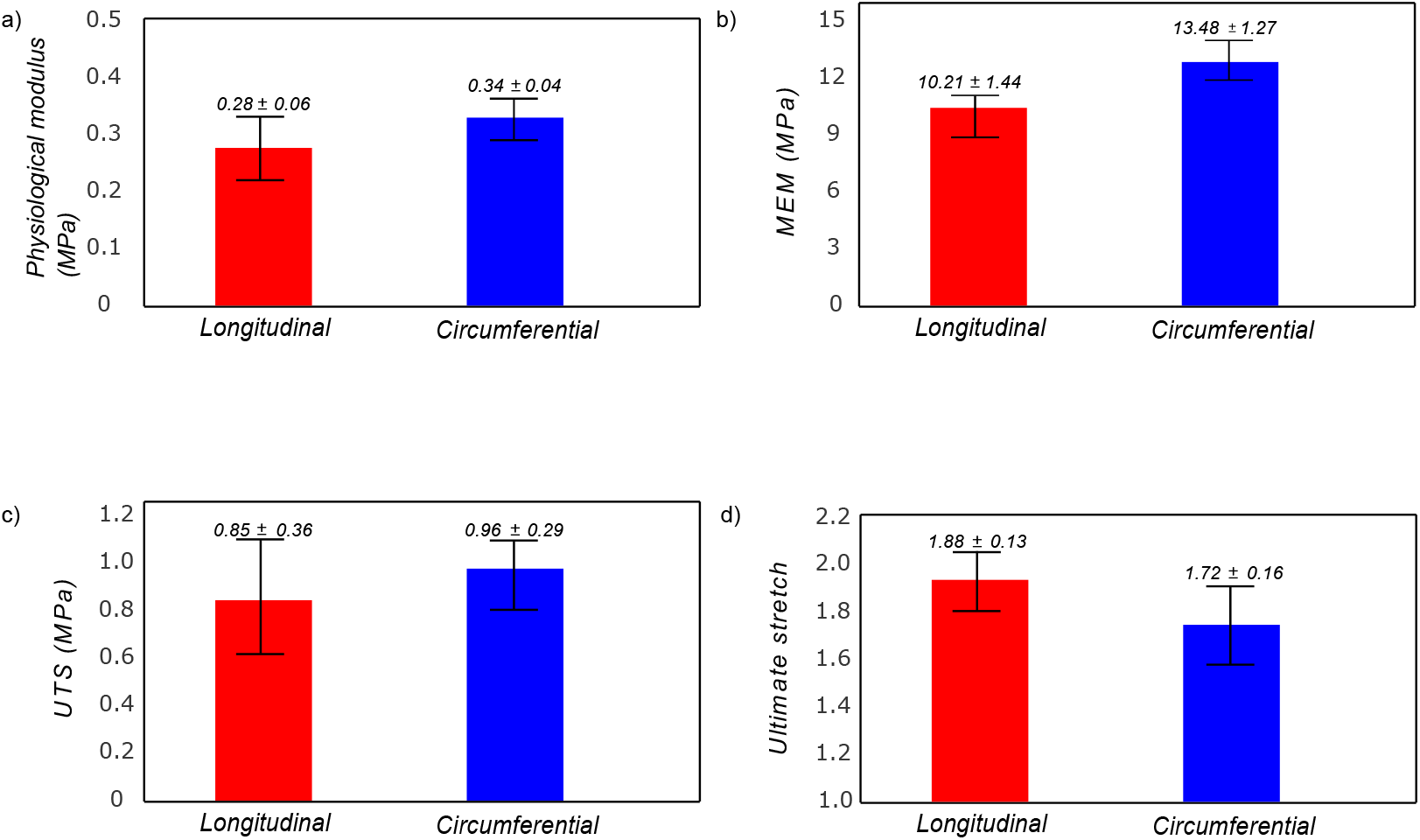
Descriptive statistics of the porcine adventitial specimens presented as mean ± standard deviation.

**Figure 7:**
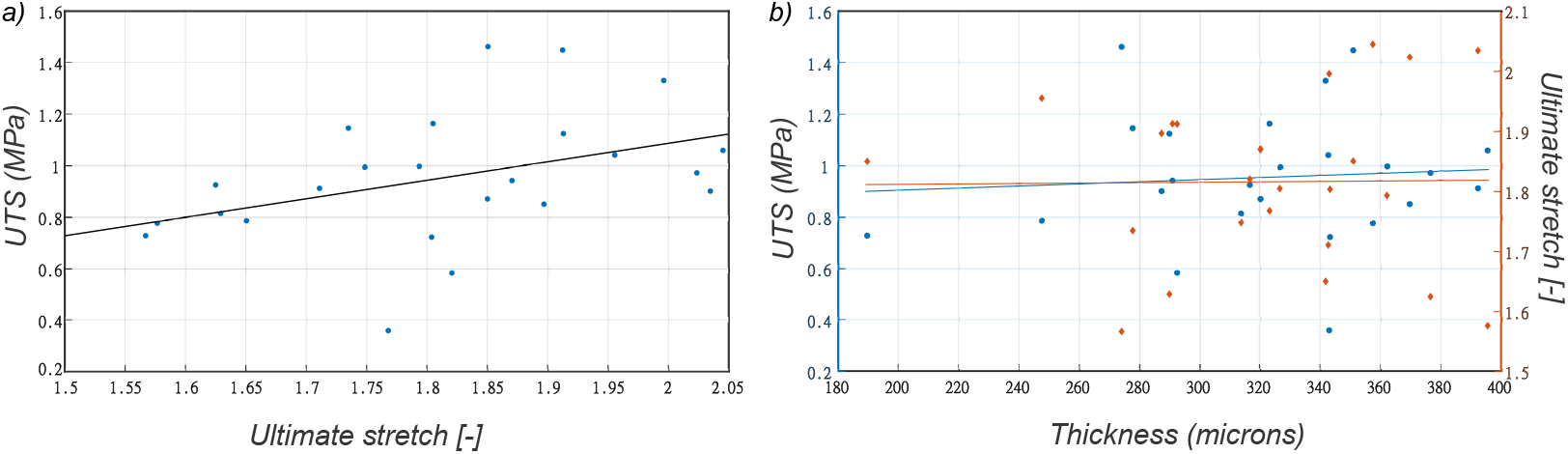
a) Correlation between measured tissue’s ultimate stress vs stretch, b) Average thickness does not correlate well with either ultimate tensile stress or ultimate tensile stretch.

**Table 1:**
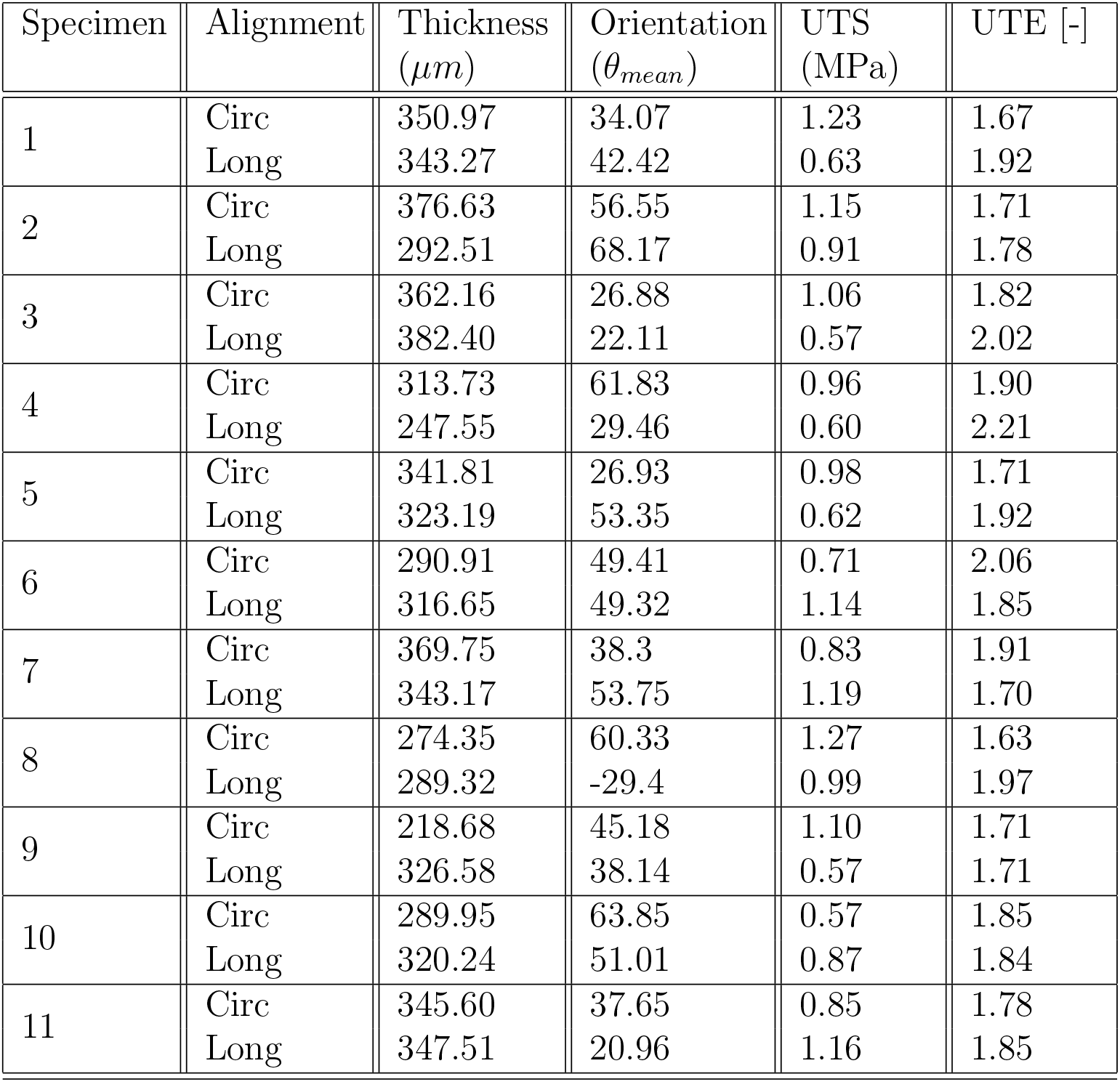
Computed parameters for the adventitial layer of all samples.

All tissue samples exhibit a pronounced nonlinear and anisotropic behavior for both elastic and rupture response. In addition to failure characteristics, the circumferential specimens displayed a higher apparent elastic modulus prior to rupture (13.48 ± 1.27 *MPa*) as compared to the longitudinal specimens (10.21 ± 1.44 *MPa*). The stress versus stretch plots for the specimens of the two groups are depicted in figure 5, but the failure stress (*UTS*) revealed no significant correlation with the failure stretch (*Ultimatestretch*), as shown in figure 6a. It was found that the adventitial thickness did not correlate with either failure stress or failure stretch either as shown in figure 6b.

### 3.2. Material modeling

Generally, arterial tissue is modelled as an anisotropic hyperelastic material [10]. In this study, an anisotropic hyperelastic constitutive model with fiber-level elasto-plastic damage proposed by Holzapfel [40] is adopted for the strain energy function. The identified optimal material parameters for all specimens are presented in table 2. A representative comparison of experimental result to the model response performed on specimen VII is shown in figure 6. On average the model prediction of both the elastic and failure domain of the mechanical response agreed well with the experimental observations (*r*^2^ = 0.94±0.04). The average identified isotropic material constant for circumferential and longitudinal specimens is 28.7±3.4 *kPa* and 35.6±8.1 *kPa* respectively. The average identified anisotropic material parameters for circumferential and longitudinal specimens are *k*_1*c*_ = 426.6 ± 107.3 *kPa, k*_2*c*_ = 6.3 ± 2.8; and *k*_1*l*_ = 568.2 ± 129.4 *kPa, k*_2*l*_ = 5.1 ± 3.4 respectively. Lastly, identified critical fiber stretch for circumferential and longitudinal specimens are *λ_c_* = 1.19 ± 0.07 and *λ_c_* = 1.24 ± 0.05 respectively. The two groups of identified material parameters were not significantly different (*p* – *value* < 0.05).

### 3.3. Structure vs tissue failure

The average orientation of the fibers with respect to the loading direction for circumferential and longitudinal specimens was observed to be 40.4±13.2 degrees and 46.8 ± 16.3 degrees respectively. Figure 9.a and 9.b depict the global failure stress (UTS) and the global failure stretch (ultimate stretch) values in relation to the mean fiber angle measured relative to the loading direction. Both the measured global failure stress and global failure stretch show a dependence on the mean fiber orientation. From figure 9 it is evident that both the global stress and stretch values decrease with increasing fiber orientation away from the loading direction.

**Figure 8:**
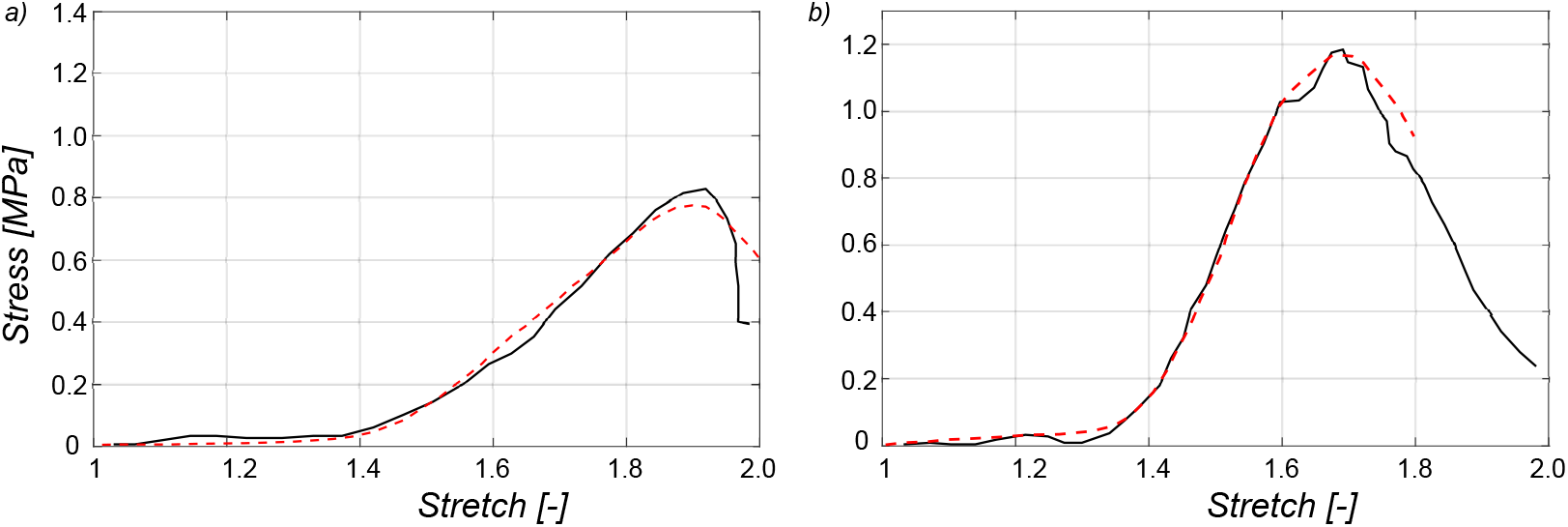
Evaluation of the proposed elasto-plastic damage model for tissue failure shows good comparison to experimental curves: a) cricumferemtial samples, b) longitudinal sample.

**Figure 9:**
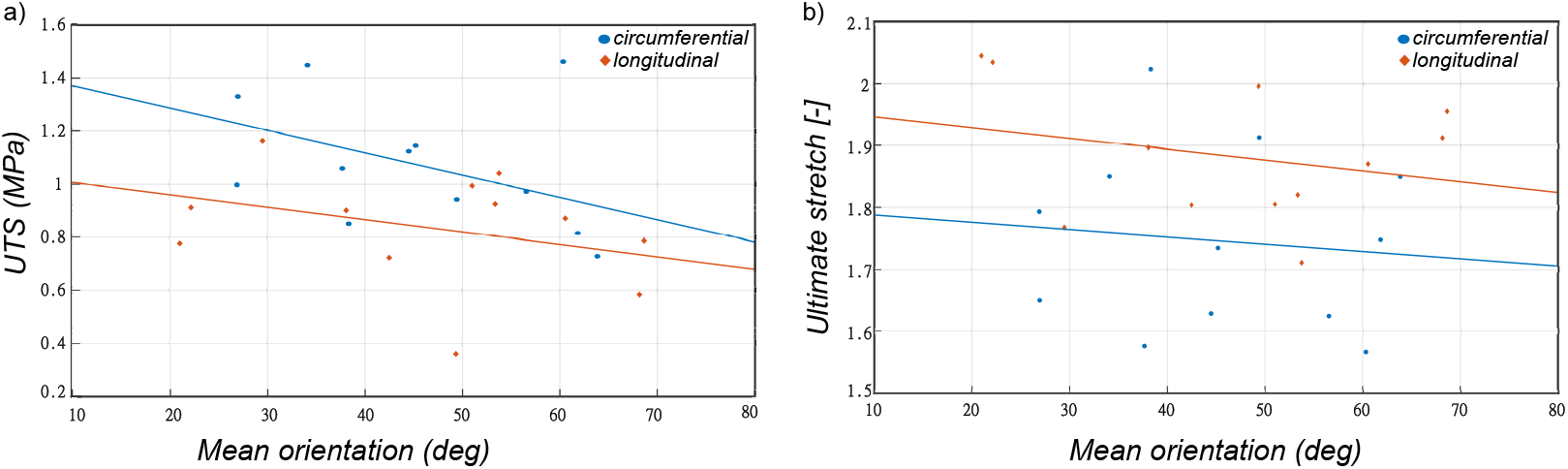
Correlation between fiber morphology and tissue’s failure properties: a) fiber mean orientation vs tissue’s ultimate tensile stress, b) fiber mean orientation vs tissue’s ultimate tensile stretch.

## 4. Discussion

The arterial tissue is a complex hierarchical structure whose functionality can be impaired by several cardiovascular diseases that cause damage, such as aneurysm or dissection, leading to a high risk of mortality. Given the highly undulated structure of adventitial collagen, combined with their anisotropic organization, it is often accepted that the adventitia contributes to the arterial wall’s mechanical properties at higher strains only [43, 44]. Experimental analyses showed that in in-vivo state, the medial layer carries up to 98% of the stress [45]. This supports both the hypotheses that media carries the physiological load and that adventitia is optimized to protect media at supra-physiological loads or in the event of disease. In that context, the purpose of this study was to provide a better understanding of the possible micro-scale phenomena at the level of collagen fibers, that lead the tissue to rupture. Through this study, we have presented a unique set of data relating to the uniaxial failure properties of porcine adventitial specimens and the underlying collagen microstructure. Layer-specific constitutive modeling of the arterial tissue has been performed for capturing both the elastic [46, 47] and failure response [26]. However, especially in the case of modeling arterial failure, a phenomenological model for anisotropic damage is limiting. In this study, we implemented an elastoplastic formulation of collagen fiber damage based on observations made during uniaxial tension experiments and reported studies on tendons [48, 49].

Nevertheless, such an error in layer separation is not expected to significantly influence the result [50].

With regard to the recorded failure stress from uniaxial tests of porcine aortic adventitia, the circumferential specimens (0.96 ± 0.29 *MPa*) showed a higher value compared to longitudinal specimens (0.86 ± 0.35 *MPa*). Layer specific failure tests conducted on ascending thoracic aneurysmatic aorta [51], healthy abdominal and aneurysmatic aorta [27], and thoracic and abdominal porcine aorta layers [26] revealed a similar trend where the circumferential direction displayed a higher rupture strength compared to the longitudinal direction. Studies conducted on the whole intact arterial wall also reported a similar trend of higher failure stresses in the circumferential than in the longitudinal direction [52, 53]. However, the study conducted by Vorp et al [54] found almost isotropic failure properties of thoracic aortic aneurysms with similar mean failure stresses in the circumferential and longitudinal directions (1180 versus 1210 kPa). Most studies observed anisotropic failure stresses for healthy and aneurysmatic human thoracic aortic medias as well as for the intact wall, with higher stresses in the circumferential than in the longitudinal direction. The anisotropic failure stresses can often be be explained by the intrinsic fiber assembly of the tissue.

However, the mean failure stretch was found to be higher in the longitudinal specimens (1.88 ± 0.13) compared to circumferential specimens (1.72 ± 0.16). Pena et al [26] reported a similar trend in both descending thoracic and abdominal aortic adventitial strips with circumferential to longitudinal failure stretches ratios of 0.93 and 0.84 respectively. Similarly, Teng et al [27] et al reported greater extensibility in the longitudinal direction in both normal and aneurysmatic aortas. A few studies [51, 55], however, reported a higher extensibility in the circumferential direction. Likewise, for the whole intact wall, failure stretches were found to be the same in both directions for the intact wall [53], but also larger in the circumferential than in the longitudinal direction [56].

The overall mean fiber orientation with respect to loading direction was found to be similar in both circumferential (40.4*°* ± 13.2*°*) and longitudinal specimens (46.8*°* ± 16.3*°*). A mean orientation close to the diagonal direction (±45*°*) was also reported for porcine aortic adventitia [30].

With regard to figure 9.a, we see observe that the ultimate stress decreases with the mean fiber orientation with respect to the loading direction. The coefficient of correlation was found to be *r* = −0.36 for the circumferential specimens and *r* = −0.28 for the longitudinal specimens. Specifically, the higher the mean fiber angle from the loading direction the lower the failure stress. Further investigation is needed to clarify the possible influence of fiber dispersion (both in-plane and out of plane) and waviness. With regard to figure 9.b, both specimens showed a weak negative correlation between ultimate stretch and mean fiber orientation. The computed coefficient of correlations are *r* = −0.11 and *r* = −0.17 for the circumferential and longitudinal specimens respectively. This may be in part explained by the waviness of the collagen fibers in the unloaded state.

Several damage mechanisms have been proposed for predicting arterial tissue rupture, ranging from tissue scale models [57] to molecular level damage models [58]. However, the majority of the models employed are phenomenological. Fibrillar slippage and kinematic disorders are possible micro-scale damage mechanisms reported on highly-organized collagenous structures such as tendons and ligaments [49]. Following that, a fiber scale elasto-plastic damage model is applied in this study to predict tissue scale rupture. The model assumes a critical stretch for individual collagen fibers for initial of damage. The identified critical stretch for initiation of fiber damage was 1.20 ± 0.06 in the circumferential specimens and 1.23 ± 0.05 in the longitudinal specimens. The study conducted by Miyazaki and Hayashi on individual collagen fibers of diameter 1 – 10 μm showed a failure stretch of 1.216±0.03 [59]. Similarly, the study conducted by Gentleman et al [60] revealed a fiber rupture stretch of 1.20 – 1.25. Although the specimens investigated in this study exhibited a wide range of fiber failure stretches, they are still within an acceptable range of experimental observations. This is implicative of fiber de-bonding and rupture being probable micro-scale damage mechanisms leading up to rupture in arterial tissue. Further investigations by combining micro-structural imaging and mechanical testing of the tissue (in multiple loading scenarios) for diseased aneurysmal tissues are imperative.

With regard to figure 10.a, the identified critical fiber stretch displayed a weak positive correlation with the experimentally recorded ultimate tensile stress. The computed coefficient of correlations are *r* = 0.18 and *r* = 0.12 for the circumferential and longitudinal specimens respectively. However, as shown in figure 10.b, the critical fiber stretch had a stronger positive correlation with the experimentally recorded ultimate stretch. The computed coefficient of correlations are *r* = 0.58 and *r* = 0.70 for the circumferential and longitudinal specimens respectively. The discrepancy in both trends is poorly understood at the moment. Further, the huge variation in the identified critical fiber stretch indicates multiple possible failure mechanisms working simultaneously towards tissue scale failure.

**Figure 10:**
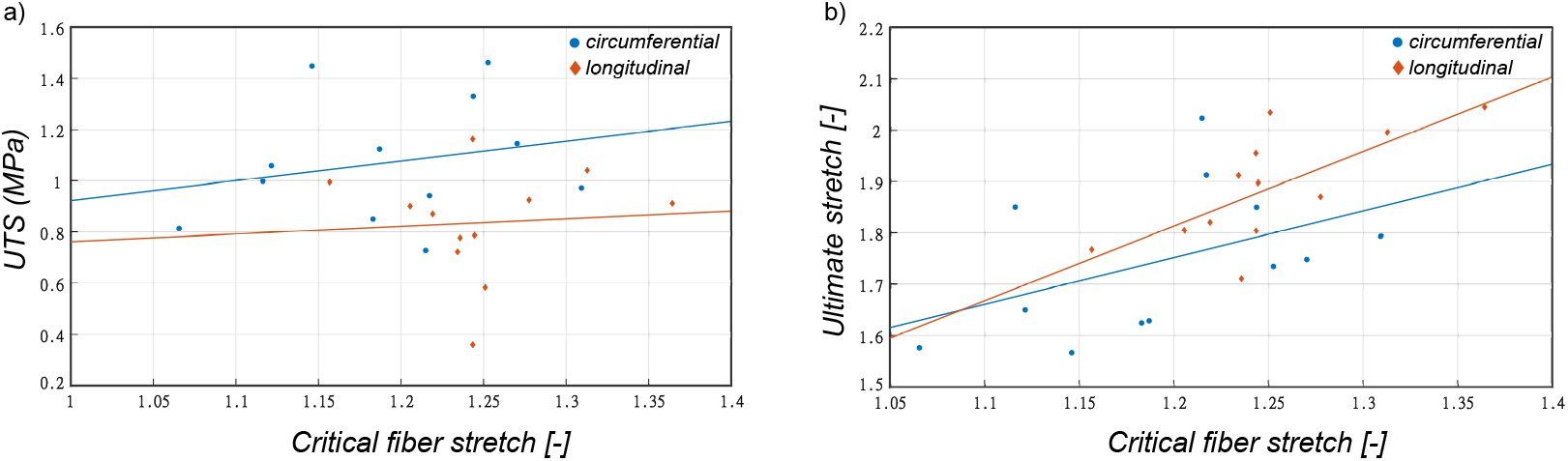
Correlation plots between critical fiber failure stretch and a)tissue’s ultimate tensile stress, b)tissue’s ultimate tensile stretch.

There are a number of assumptions and limitations in the current work that should be addressed:(1) uniaxial tension tests were chosen to achieve failure of the tissue, which is significantly difficult under planar biaxial tension. Hence, this is insufficient to capture the anisotropic material behaviour of the tissue. Moreover given that the in-vivo loading is close to biaxial, the ultimate tensile strength and ultimate stretch obtained in this study may be different from the true values; (2) it was assumed that the samples were in a stress-free state at the beginning of each test but the specimens were not perfectly flat at the beginning of the test and this led to some small initial deformation; (3) in this study, an assumption was made that the deformation is the same throughout the whole thickness; (4) optical microscopy was used as a simplified imaging tool in place of multi-photon microscopy to characterize the collagen fiber morphology. Although this provides a larger field of view, it suppresses information through the thickness, thus only providing an aggregate assessment; (5) only the mean orientation of the fibers was considered in the damage constitutive model. Other structural parameters such as in-plane dispersion, out of plane dispersion, fiber waviness, and content are all important in interpreting the micro-scale mechanisms of tissue failure; (6) The constitutive parameters were estimated by fitting individually the stress–stretch relationships resulting from the uniaxial tensile testing in the circumferential and axial directions; (7) it is assumed that the collagen density is same for the all images, which is a strong simplification. Measuring collagen density through quantitative histology would improve the accuracy of identified stiffness related material parameters; (8) even though there is elastin present in the adventitia, it was assumed that the isotropic material parameter includes the contribution of elastin fibers along with the matrix.

## 5. Conclusion

The present study provides data on the direction specific failure properties of healthy porcine abdominal aortic adventitial tissue. Combining failure tests with mechanical testing highlights the strong influence of the collagen microstructure on its failure properties. Further, in this study, we quantify a novel fiber-level approach for damage initiation and ultimately failure. Combining uniaxial failure mechanical data with a structural constitutive model that takes into account the mean fiber orientation in the tissue enabled us to quantify meaningful micro-scale damage mechanisms. The study helps in expanding the the research on developing predictive capabilities of constitutive models fed through thorough experimental investigations. This could be further used for predicting the failure properties of soft biological tissues, and is a step towards realistic modeling of tissue failure. Finally, this would help in addressing clinical questions regarding rupture of aneurysmatic aortas with appropriate medical imaging techniques.

## 6. Acknowledgements

This work was funded by the European Research Counsil, starting grant no.638804, AArteMIS.

